# Molecular evolution of silver nanoparticle resistance in a bacterial pathogen and a unique adaptation response to ionic silver

**DOI:** 10.1101/2025.01.28.635224

**Authors:** Oliver McNeilly, Daniel G. Mediati, Matthew J. Pittorino, Riti Mann, Bill Söderström, Mehrad Hamidian, Cindy Gunawan

## Abstract

This research explores the adaptive defense mechanisms of a nanosilver-resistant pathogen (NAg^R^) to protect and fight off the complex antimicrobial targeting of silver nanoparticles. The Gram-negative bacterium *Acinetobacter baumannii* upregulated expression of outer membrane proteins for cell surface defense, as well as membrane and capsule synthesis genes. Increased abundance of surface-attached biofilm colonies in NAg^R^ was linked to the phenotypically indicated increase in membrane integrity, with the bacterium also forming more EPS, the biopolymer matrix that protects the residing colony. In response to the known reactive oxygen species (ROS) toxicity characteristics of the nanoparticle, NAg^R^ upregulated its oxidative stress management system, specifically involving ROS scavenger enzymes and opportunistic metal efflux pumps. Many of these evolved defense mechanisms only manifested in the resistant bacterium, while they were absent in the wild-type strain. This study also details the unique defenses of an ionic silver-adapted *A. baumannii* variant, having evolved from the same wild-type parental strain as NAg^R^. Despite similarities in cell surface and biofilm defense trends, the slower-to-kill tolerant strain (Ag^T^) exclusively upregulated multidrug efflux systems and respiratory chain enzymes, thought to maintain enhanced respiration activity, a known tolerant characteristic. Identification of these stable defense mechanisms can recommend strategies for molecular targeting to overcome the adaptation phenomena.

## Introduction

The medical and commercial indiscriminate use of antimicrobial agents, along with diminishing pharmaceutical investment toward novel antibiotic development, have been the primary drivers of global antimicrobial resistance (AMR)^1–3^. Between 2019 – 2021, systematic analyses estimated over 4 million deaths were attributable to AMR globally^4^. One of the major contributors to the AMR crisis is the opportunistic Gram-negative coccobacillus bacterium *Acinetobacter baumannii*. Carbapenem-resistant *A. baumannii* has been classified as the number one critical level priority pathogen by the World Health Organization since 2017^5,6^. This species is among the leading cause of nosocomial (hospital-acquired) infection, with for instance, 80 – 97% of reported cases in Southern Europe, the Middle East, and North Africa, due to multidrug-resistant (MDR) strains^7,8^. Global mortality rates from MDR *A. baumannii* have ranged from 24 – 83%, with higher incidences occurring in elderly and/or immunocompromised individuals^9–11^. Treatment options for MDR (including carbapenem-resistant) *A. baumannii* infections are currently limited to antibiotics such as polymyxins (*i.e.* colistin), and yet, resistance to these last line treatments have been identified globally^12–14^.

The growing threat of AMR calls for immediate and innovative solutions. Nanotechnology has gained significant traction in the biomedical sector in recent years^15^. Silver nanoparticles (NAg) are now one of the most commercialized products of nanotechnology, due to their unique physicochemical characteristics and broad-spectrum antimicrobial properties. The nanoparticles have been shown to be equally effective against MDR pathogens, including MDR *A. baumannii*^16–19^. Currently a major alternative antimicrobial, NAg have been incorporated in medical devices, such as in wound dressings and catheters, as well as, increasingly, in consumer products, for instance in household appliances, with the primary aims to manage pathogen growth^20–22^. NAg target microbes through the cell-killing activities of the soluble silver that leach from the nanoparticles, including ionic silver (Ag^+^), and the solid silver particulates that remain after leaching^23–25^. The antimicrobial activities have been largely associated with cellular generation of reactive oxygen species (ROS)^24,26^. Among other potential ROS-generation pathways, studies have indicated initial nanosilver targeting, as well as leached Ag^+^, of membrane-bound respiratory chain components in bacteria^24,27^. The resulting disruption of the electron transport chain causes premature leakage of electrons, thereby reducing molecular oxygen (O_2_), present in the cytoplasm, into superoxide radicals (O_2_^•-^)^24^. Studies have further suggested that O_2_^•-^ radicals (and leached Ag^+^) target iron-sulfur (Fe-S) clusters that are present in many important enzymes and proteins^28,29^. Notably, the intracellular presence of leached Ag^+^ has also been linked to potential Trojan-horse type toxicity of NAg, with intracellular ion leaching following particle uptake^30^. The targeting of Fe-S clusters could consequently release Fenton-active Fe(II) ions, which can react with cellular H_2_O_2_ to generate the highly reactive hydroxyl radicals (^•^OH)^24,29^. Hydroxyl and superoxide radicals have been shown to inactivate proteins, for instance, by oxidizing the sulfur atom in the amino acid methionine to sulfoxide and/or oxidizing the thiol group (-SH) in cysteine to thiyl radical (RS^•^)^29,31^. In DNA, hydroxyl radical targets H-bonds that are present in the nucleotide base-pairs, as well as in the sugar moieties of the sugar-phosphate backbone, causing nucleotide cleavage^32^. The radical is also well recognized for its lipid peroxidation activities, including membrane phospholipids^33^.

The increasing and often indiscriminate use of nanosilver has raised concerns over the development of silver-adapted pathogens, despite complex, multi-targeting antimicrobial activities of silver. Resistance to ionic silver have been reported in bacteria for decades, including in *A. baumannii*^34–38^. These resistance traits could develop endogenously, for instance through gene mutations, while in other cases can be acquired through uptake of mobile genetic elements, such as *sil* gene-harboring plasmids. For the latter, studies have detected presence of members of the nine silver efflux *silCFBA(ORF105aa)PRSE* genes in *A. baumannii* plasmids^39^. Evidence of resistance to nanosilver was first reported in 2013 in environmental bacteria^40^. The bacterial population was able to grow at an otherwise lethal concentration of the nanoparticle following prolonged exposure. Indeed, such adaptive resistance phenomena to nanosilver have been increasingly observed, in both Gram-negative and Gram-positive bacterial species, including clinically-relevant strains^39,41–44^. Recently, we reported the first known evidence of evolutionary-adapted nanosilver resistance in *A. baumannii* (model strain ATCC 19606), following long-term exposure to the nanoparticle^45^. With no prevalence of any of the *sil* genes, either in its chromosome or native plasmids, the bacterium was able to evolve harder-to-kill resistance traits, which saw growth at up to 5-fold nanosilver MIC (minimum inhibition concentration, of the wild-type strain). The adaptation characteristics still manifested even after growth in antimicrobial-free conditions. Rationally, such stable mechanisms motivate further in-depth work to identify the evolved defense characteristics at the molecular level.

Herein, we studied changes in gene expression and phenotypic (observable traits) profiles of the nanosilver-resistant *A. baumannii* strain (NAg^R^), in response to sub-lethal and lethal nanosilver concentrations. When compared to those exhibited in the wild-type strain (WT), it is evident that the resistant bacterium has evolved primary defense mechanisms, which were already observed in WT, and further upregulated in NAg^R^, as well as, secondary mechanisms, the latter were only seen in the resistant strain. Apart from the anticipated defense mechanisms to control the ROS-associated toxicity, the studies revealed stable silver-evolved cell surface defense mechanisms and changes in biofilm growth behavior in the Gram-negative pathogen. The present work also studied the gene expression and phenotypic profiles of an ionic silver-adaptive *A. baumannii*. The slower-to-kill tolerant strain (Ag^T^) was developed from the same ATCC 19606 parent, subjected to comparable (30-day) long-term exposures, to ionic silver^45^. Despite sharing a number of silver-relevant defense traits, Ag^T^ is physiologically distinct, having also evolved exclusive Ag^+^-protective mechanisms. Insights into the molecular basis of silver resistance can provide indication of molecular targets to help overcome the adaptation phenomena.

## Materials and Methods

### Nanosilver and ionic silver

The silver nanoparticles (6.6wt% Ag_2_O [*d*_TEM_ ≈ 2 nm] finely dispersed onto inert TiO_2_ [*d*_TEM_ ≈ 30 nm]) were synthesized *via* flame spray pyrolysis, as described by Gunawan *et al.*^40^. The nanoparticles were sterilized with gamma-irradiation (Cobalt-60) for 1 h at ∼ 6 Gy/min dose at the Australian Nuclear Science Technology Organization (ANSTO). A fresh stock of NAg suspension was prepared in sterile cation-adjusted Mueller-Hinton broth (CAMHB; BD) and homogenized *via* ultra-sonication (20 s, 50% output; Vibra-cell, Sonics & Materials), prior to studies. Ionic silver was supplied as silver nitrate (AgNO_3_; Merck) suspended in sterile ultrapure Milli-Q water. All NAg and Ag^+^ exposure experiments were performed in dark conditions, to photo-catalytically inactivate the TiO_2_ support, and to prevent reduction of Ag^+^ to Ag^0^, respectively^25,40^.

### Bacterial strains and growth conditions

The wild-type (WT) *A. baumannii* ATCC 19606 strain used in this study was first isolated from a patient urine sample in 1948 by Schaub and Hauber^46^. The NAg-resistant (NAg^R^) and Ag^+^-tolerant (Ag^T^) strains were developed from ATCC 19606 through 30-day exposures to NAg and Ag^+^, respectively, in our previous study^45^. Frozen (−80°C) glycerol stocks of the WT, NAg^R^, and Ag^T^ strains were streaked onto cation-adjusted Mueller-Hinton agar (CAMHA) plates and grown for 16-18 h at 37°C. For all experiments, single isolated agar colonies were inoculated in CAMHB and grown overnight for 16-18 h at 37°C, 250 rpm.

### Working concentrations of NAg and Ag^+^

To ensure minimal compensatory effects (as fitness cost trade-offs)^47^, we compared the early exponential phase growth rates of the WT, NAg^R^ and Ag^T^ strains with and without the NAg and/or Ag^+^ working concentrations. Briefly, overnight cultures of each strain were diluted in fresh CAMHB to OD_600_ 0.05, then cultured at 37°C, 250 rpm for 2 h. The cultures were then exposed to the low (0.5 x MIC, 0.5 µg Ag/mL for NAg, 1 µg Ag/mL for Ag^+^)^45^ and high (3 µg Ag/mL for NAg (3 x MIC) and Ag^+^ (1.5 x MIC))^45^ silver doses (**Table S1**), with hourly OD_600_ readings. Note that the different MIC-fold for the high NAg and Ag^+^ exposure concentrations were to ensure equivalent silver doses (3 µg Ag/mL). The growth profiles are shown in **Figure S3 to S8**, with comparable early exponential phase growth rates of the silver-exposed strains relative to their respective cell-only (no silver) growth, hence indicating a silver-only induced effects. From herein after, the mRNA and phenotypic studies were performed at the above-mentioned silver concentrations. The growth profile studies were performed with at least two biological replicates, each with technical replicates.

### RNA (transcriptomic) studies

The low and high concentration silver-exposed and cell-only cultures of WT, NAg^R^ and Ag^T^ strains were harvested at 30 mins of exposure, or growth at 37°C, in the exponential phase (silver was added after 2 h growth, see above). The 30-min exposure timeframe allowed direct profiling of the immediate silver-induced defense responses prior to the known extensive cell-killing activities at 1 hour of exposure (at MIC level)^23,24^. See **Figure S3 to S8** for the OD_600_ readings of the silver-exposed and cell-only cultures at the time of RNA isolation.

#### RNA extraction and sequencing

Total RNA extraction was performed using the RNeasy Mini Bacteria Kit (Qiagen) with RNAprotect Bacteria Reagent (Qiagen) following manufacturer’s instructions. Briefly, 0.5 mL of each of the silver-exposed (and cell-only) cultures was incubated with 1 mL RNAprotect (5 min, room temperature) and pelleted down (7500 rpm, 10 min). Cell pellets were lysed with lysozyme (Qiagen), with addition of Proteinase K (20 µL 10 mg/mL, Sigma-Aldrich), the latter to digest proteins and remove nucleases, for 15-20 min, room temperature (10 s vortex every 2 min), followed by DNase treatment for 15 min, room temperature. RNA extracts were then eluted from the spin column with RNase-free water (13000 rpm, 1 min), followed by a dry spin. RNA quantity and purity was assessed using an Epoch microplate spectrophotometer (Agilent Technologies) and a Nanodrop spectrophotometer (Thermo-Fisher Scientific). RNA integrity was assessed using the TapeStation system with RNA Analysis ScreenTape (Agilent Technologies). All purified RNA samples contained RIN values of > 8 (data not shown). Ribosomal RNA depletion, library preparation, and sequencing were performed at the Ramaciotti Centre for Genomics (UNSW, Australia) on a NovaSeq6000 platform (Illumina) generating up to 800 million single-end 100 bp reads.

#### RNA data analysis

RNA-seq datasets (chromosomal and native plasmid transcripts) were analyzed using the limma-voom software^48^, in Galaxy Australia v3.58.1 (https://usegalaxy.org.au). The analysis pipelines, including quality control of reads *via* multi-dimensional principal component analysis (see **Figure S2**), read coverage, counts and mapping, identification of differentially expressed genes (DEGs), and gene ontology clustering, are shown in **Figure S1**, each with the software used. Briefly, the demultiplexed sequence reads underwent quality control analysis with FastQC. Trimmed and filtered reads (using Trimmomatic, allowing a minimum of 40-nt sequence) were aligned to the ATCC 19606 chromosome (GenBank accession number CP045110) and the two ATCC 19606 native plasmids, p1ATCC19606 and p2ATCC19606 (GenBank acc. no. CP045108 and CP045109) using the read mapper Bowtie2, allowing for one mismatch^49^. The frequency of mapped reads in each BAM file that aligned to the “gene” or “CDS” feature of the ATCC 19606 GFF3 file was calculated and used to generate tabular datasets for each sample using the count quantification software HTseq. The tabular datasets were used to identify DEGs between biological replicates and treatment conditions. The cut-off threshold to define a statistically significant DEG between pairwise comparisons was set at a log_2_-fold change of ≥ 0.58 (≥ 1.5-fold change) with an adjusted *p*-value of ≤ 0.01, as per the Benjamini and Hochberg false-discovery rate (FDR) method^50^. Mapping of statistically significant DEGs to functional pathways with gene ontology was performed using ShinyGO (http://bioinformatics.sdstate.edu/go/) and the results are reported in **Table S2 and Table S3**. Individual DEGs of interest associated with silver defense mechanisms of the WT, NAg^R^ and Ag^T^ strains were further examined. A summary outlining the silver exposure systems and details for each pairwise comparison (with total DEGs identified) are provided in **Table S4 and Table S5**.

### Data Availability Statement

The RNA-seq datasets (chromosomal and native plasmid transcripts) that support the findings of this study are openly available in NIH National Center for Biotechnology Information at https://www.ncbi.nlm.nih.gov/bioproject/PRJNA557095/, BioProject accession number PRJNA557095^124^.

### Microscopy phenotypic studies

#### Bacterial membrane study

The WT, NAg^R^, and Ag^T^ strains were grown in the absence of silver. Upon reaching the exponential growth phase (2 h growth, see above), the cells were directly stained with FM 4-64 (10 μg/mL working concentration, Thermo-Fisher Scientific) for 10 min in the dark at room temperature. The stained cells were then pipetted onto 1.5% (w/v) agarose gel pads and imaged using DeltaVision (DV) Elite deconvolution fluorescence microscope (GE Healthcare), with mCherry filter (572/25 nm excitation, 632/60 nm emission) to visualise stained cell membranes. The images were analysed with the Fiji (ImageJ, USA) plugin MicrobeJ^51^.

#### Biofilm study

For biofilm growth, overnight cultures of WT, NAg^R^ and Ag^T^ strains were diluted to an OD_600_ 0.05 in fresh CAMHB, inoculated into a FluoroDish (World Precision Instruments) and grown at 37°C for 24 h in humidified conditions. Following removal of the supernatant, the surface-attached biofilms were washed twice with 1x phosphate-buffered saline (PBS). For silver exposures, the biofilms were treated with their respective silver agents (NAg and/or Ag^+^, 0.5 x MIC), for 24 h at 37°C^45^. Untreated cultures for each strain were included. The cultures were washed twice with PBS, the biofilms were then stained with SYTO-9 (5 μM working concentration, Thermo-Fisher Scientific) and FilmTracer SYPRO Ruby (1x concentration per manufacture instructions, Thermo-Fisher Scientific) for 30 min at room temperature under dark condition. The stained biofilms were washed twice with PBS for imaging.

Fluorescence imaging was performed using the DV Elite fluorescence microscope with FITC filter (475/28 nm excitation, 523/48 nm emission) for SYTO-9 (stained cells) and TRITC filter (632/22 nm excitation, 679/34 nm emission) for SYPRO Ruby (stained EPS/matrix proteins). All images were captured in sectional Z-stacks using the DV Elite SoftWoRx program and deconvolved using proprietary settings. The deconvolved biofilm images were analyzed using Imaris software v9.6.0 (Oxford Instruments) to determine colony biomass and EPS biomass (μm^3^/μm^2^) with automatic threshold (absolute intensity) detection applied. The 3D Z-stack images were compressed and presented as single plane images to provide visualisation of the biofilm structures.

#### Cellular ROS study

The WT, NAg^R^ and Ag^T^ strains were first grown for 2 h (see above). The cells were pelleted, washed twice with PBS, re-suspended in sterile saline (8 g/L NaCl, 0.2 g/L KCl), and stained with ROS-reporter 2′,7′-dichlorodihydrofluorescein diacetate (H_2_DCFDA, 10 µM working concentration, Invitrogen) for 45 min at room temperature under dark conditions. The stained cultures were then pelleted, re-suspended in saline, and exposed to silver (NAg and/or Ag^+^, 0.5 x MIC) for 30 min and 60 min at 37°C. Finally, the cultures were re-pelleted and re-suspended in saline. For imaging, the cells were pipetted into Gene Frame (2% (w/v) agarose gel pad, Thermo Fisher Scientific). Cells treated with 50 mM hydrogen peroxide (H_2_O_2_) were used as positive control.

Fluorescence imaging was performed using the DV Elite microscope with FITC filter (475/28 nm excitation, 523/48 nm emission). The images were captured using the DV Elite SoftWoRx program and deconvolved. Five-to-six images were captured per biological triplicate, per strain and silver exposure system. For fluorescence quantitative analysis, background noise was subtracted with a rolling ball radius of 50 pixels and intensity values were calculated from individual cells using the Fiji (ImageJ) plugin MicrobeJ^51^.

## Results and Discussion

### Cell surface defense and physiological changes

Herein, we first described the first-line of defense to nanosilver antimicrobial targeting in the bacterium; more specifically, on the transcriptomic and phenotypic changes related to cell envelope components which evolved as a result of long-term exposure. The wild-type (WT) strain exhibited increased expressions of genes that encode outer membrane-embedded proteins upon exposures to NAg (0.5 x MIC, 0.5 µg Ag/mL, referred to as ‘low’ nanoparticle concentration). Firstly, *oprC*, encoding the TonB-dependent copper channel OprC, was upregulated by ∼18-fold, relative to the cell-only (no silver) control samples (**Figure 1A**, *p*-adjusted ≤ 0.01). Increased OprC expression in *A. baumannii* has been indicated to enhance sequestration of toxic levels of Cu^+^ within the periplasmic space, with protein structure-based studies further suggesting the role of the protein to sequestrate or ‘trap’ silver particulates (and the leached Ag^+^, as with Cu^+^ - a soft acid) in the periplasm^52–56^. The NAg exposures also saw the WT strain upregulating the expressions of some of the most conserved outer membrane proteins in *A. baumannii*, with ∼1.7-1.9-fold upregulations of *ompA*, *ompW* and *carO*, relative to the cell-only samples. OmpA is the most conserved outer membrane protein in *A. baumannii*, with studies linking its upregulation to enhance membrane integrity, and in turn, higher resilience against many surface-targeting antimicrobial agents^57–61^. The outer membrane protein OmpW has also been associated with bacterial defense through improved cell membrane integrity^62^. The porin CarO is also highly conserved in *A. baumannii*^63^. Upregulation of this porin has been suggested to enhance cell-to-surface attachment and promote biofilm growth^57,64^. Indeed, studies have also linked OmpA and OmpW to biofilm formation in *A. baumannii,* which, as later shown in the present study, could contribute to NAg defense^57,63,65,66^. The upregulations of the outer membrane proteins were also observed with ionic silver exposure (Ag^+^, supplied as AgNO_3_). A ∼1.9-13-fold increased expressions were seen with *oprC*, *ompW* and *carO* in the wild-type strain when exposed to the ‘low’ ionic silver concentration (0.5 x MIC, 1 µg Ag/mL), relative to the cell-only control (**Figure 1B**).

**Figure 1.**
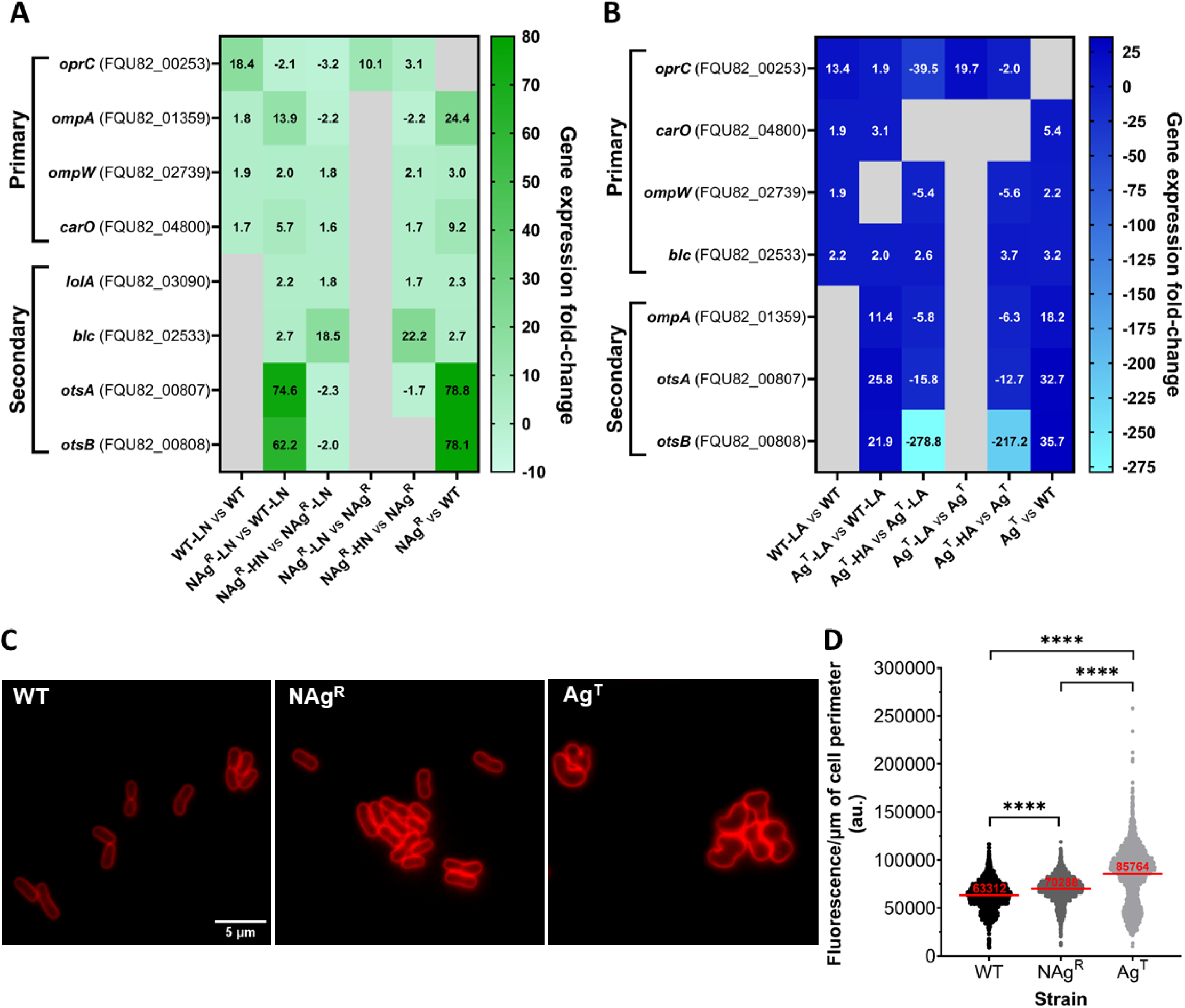
Cell surface defense mechanisms identified in silver-adaptive strains of *A. baumannii*. **(A,B)** Comparative analysis of chromosomal mRNA transcript levels of outer membrane protein (*oprC*, *ompA*, *ompW*, *carO*), membrane synthesis (*lolA*, *blc*) and capsule synthesis (*otsA*, *otsB*) genes in wild-type (WT), nanosilver-resistant (NAg^R^) and ionic silver-tolerant (Ag^T^) strains of *A. baumannii*, upon exposures to the ‘low’ silver concentrations (0.5 µg Ag/mL for nanosilver (LN, 0.5 x MIC), 1 µg Ag/mL for ionic silver (LA, 0.5 x MIC)) and ‘high’ silver concentrations (3 µg Ag/mL for nanosilver (HN, 3 x MIC) and ionic silver (HA)). The differentially expressed genes (log_2_-fold change cut-off threshold ≥ 0.58 (≥ 1.5-fold change), *p*-adjusted ≤ 0.01, grey cells indicate statistically insignificant *p*-adj > 0.01 changes) were captured at 30 min of silver exposures in the exponential growth phase Also shown are the physiological gene expression changes when in the absence of silver. (**C**) Phenotypic studies of bacterial membrane with lipophilic fluorescence staining of WT and the silver-adaptive NAg^R^ and Ag^T^ strains (scale bar = 5 µm) at 30 min growth in the exponential phase. (**D**) Quantitative analysis of fluorescence intensity per µm of cell perimeter (au.). Each data point represents a single cell measurement, with the horizontal bar showing the mean fluorescence from 2000 – 4000 cells analyzed, per strain. Statistical analysis (unpaired *t*-test with Welch’s correction, to account for unequal variances and large data sets) showed statistically significant differences in the fluorescence intensity between strains (**** *p* < 0.0001). The mRNA and phenotypic work were performed with a minimum of three biological replicates (independent bacterial inocula from different colony isolates).

These membrane proteins were upregulated at even higher levels in the resistant strains. The nanosilver-resistant strain (NAg^R^) increased the expressions of *ompA*, *ompW* and *carO* by ∼2.0-14-fold, relative to WT, when exposed to the low nanoparticle concentration (0.5 x MIC) (**Figure 1A**). Exposures to ‘high’ nanoparticle concentration (3 x MIC, 3 µg Ag/mL) saw further ∼1.6-1.8-fold upregulations of *ompW* and *carO* in the resistant strain, relative to the low concentration exposures (0.5 x MIC). Notably, *ompA* was downregulated by ∼2.2-fold, following high concentration exposure, relative to the low concentration exposure. Comparable transcriptomic changes were also seen in the case of ionic silver-tolerant strain (Ag^T^), with ∼11- and 3.1-fold increased expressions of *ompA* and *carO*, upon exposures to the low ionic silver concentration (0.5 x MIC), relative to WT (**Figure 1B**). Downregulation of *ompA*, by ∼5.8-fold, was also observed in Ag^T^, at the high concentration exposure (3 µg Ag/mL), relative to the low concentration exposure. Downregulation of *ompA* has been previously reported in silver-exposed *E. coli*, however, the exact reason for this remains unclear^35^. Next, we also observed downregulation of *oprC*, in fact, in both NAg^R^ and Ag^T^ strains, when exposed to the low silver concentration (0.5 x MIC), by ∼2.1-fold in NAg^R^, relative to WT, and when exposed to the high silver concentration (3 µg Ag/mL), by ∼3.2-fold in NAg^R^, and by the more substantial ∼40-fold in Ag^T^, relative to the low concentration exposures. The reasons for these downregulations are still not clear, although they most likely relate to management of periplasmic silver levels^67^.

Next, we found that the nanosilver-resistant strain upregulated a number of lipoprotein carriers, which were not seen in the wild-type strain, hence hereby defined as secondary defense mechanisms (**Figure 1A**). Exposures to the high nanoparticle concentration (3 x MIC, 3 µg Ag/mL) saw ∼1.7- and ∼22-fold increased expressions of *lolA* and *blc* (lipocalin), respectively, in the NAg^R^ strain, relative to the cell-only samples. No statistically significant upregulations were observed in NAg^R^ with the low concentration nanoparticle exposures (0.5 x MIC), relative to the cell-only samples, however, a ∼2.2- and ∼2.7-fold respective upregulations were detected, relative to WT. LolA is a periplasmic chaperone that forms a part of the LolABCDE protein complex. LolA, like the outer membrane-embedded protein lipocalin, transports lipoproteins for outer membrane biogenesis^68,69^. LolA, along with the ATP-binding cassette transporter LolCDE, shuttles newly synthesized lipoproteins to the outer membrane^68^. Increased expressions of the Lol system and lipocalins, have been observed in antimicrobial exposure cases in bacteria, which are thought to increase lipoprotein delivery to the outer membrane for repair^68,69^. Relevant to nanosilver toxicity, studies have reported upregulations of lipocalins in Gram-negative bacteria, including *E. coli*, in response to oxidative stress^70–72^. Upregulations of *blc* were also observed in Ag^T^, by ∼3.7-fold with the high concentration exposures (3 µg Ag/mL), relative to the cell-only samples, and by ∼2.0-fold with the low concentration exposures (0.5 x MIC), relative to WT (**Figure 1B**).

Also considered as secondary mechanisms, are the herein observed upregulations of *otsA* and *otsB* in the resistant strain, which respectively encode the trehalose synthesis enzymes trehalose-6-phosphate synthase and trehalose-6-phosphate phosphatase^73,74^. A structural component of surface capsular polysaccharide (CPS), increased trehalose synthesis has been indicated in bacteria under stress conditions^75^. The NAg^R^ strain increased the expressions of *otsA* and *otsB* upon exposures to the low nanoparticle concentration (0.5 x MIC), at the considerably high ∼62- and 75-fold, relative to WT (**Figure 1A**). In *A. baumannii*, increased trehalose synthesis has been associated with enhanced CPS density and thickness, which is thought to contribute to cell envelope protective effects^74^. In fact, upregulations of CPS synthesis genes have been previously observed with NAg exposures, in *E. coli* and *S. aureus*^76,77^. Upregulations of *otsA* and *otsB* were also observed in Ag^T^, by ∼26- and 22-fold, relative to WT, when exposed to the low ionic silver concentration (0.5 x MIC) (**Figure 1B**). These genes, however, were downregulated at the high silver concentration exposures (3 µg Ag/mL, nanoparticle and ionic silver), by ∼2.3- and 2.0-fold, respectively, in NAg^R^, and by the more substantial ∼16- and ∼279-fold, respectively, in Ag^T^, relative to the low silver concentration exposures (0.5 x MIC). The downregulations, particularly the extensive ones, again, seen with Ag^T^, are thought to form part of the bacterium’s strategy to conserve energy for the exclusive silver defense mechanisms that only manifested in this strain, as later described.

The transcriptomic changes in the nanosilver-resistant strain were in fact, already manifested even when in the absence of silver. Upregulations of the outer membrane proteins (*ompA*, *ompW*, *carO*) by ∼3.0-24-fold, the lipoprotein carriers (*lolA* and *blc*) by ∼2.3- and 2.7-fold, respectively, and again, the notably high upregulations of the capsule synthesis enzymes (*otsA* and *otsb*) by ∼78-79-fold, were observed in the NAg^R^ strain, relative to the WT strain (no silver, **Figure 1A**). This indicates that stable cell envelope integrity (and membrane repair) defense mechanisms have evolved in the resistant strain. Detailed fluorescence analysis of NAg^R^ stained with FM4-64 showed increased membrane fluorescence (per µm of cell perimeter), when compared to WT (*p* < 0.0001, no silver, **Figure 1C, D**). The highly lipophilic dye (firstly) stains the bacterium outer membrane (before diffusing intracellularly), intercalating itself in the membrane phospholipids and lipoproteins^78,79^. In agreement with the transcriptomic changes, the increased membrane fluorescence intensity is thought to result from an increase in membrane density. These phenotypic changes were indeed also observed with the ionic silver-tolerant strain (Ag^T^), with higher membrane fluorescence intensity, relative to WT (*p* < 0.0001, no silver, **Figure 1C, D**). Upregulations of *ompA*, *ompW*, *carO* by ∼2.2 – 18-fold, *blc* by ∼3.2-fold, and more substantially, *otsA* and *otsB* by ∼33- and 36-fold, respectively, were detected in Ag^T^, relative to WT (no silver, **Figure 1B**). Despite the transcriptomic and phenotypic similarities, a more detailed analysis revealed different membrane fluorescence traits with Ag^T^ when compared to NAg^R^, which could associate with the unique morphological transformation observed with the former, as later described.

### Evolved changes in biofilm growth behavior

In line with the earlier described upregulations of outer membrane proteins, as well as membrane and capsule synthesis proteins, the present study further observed increased expressions of other biofilm growth-associated genes, evolving from the silver prolonged exposures. Bacterial community when growing as surface-attached biofilms are protected from antimicrobial targeting by matrix of polysaccharides, proteins, lipids and (extracellular) DNA, the so-called EPS (extracellular polymeric substance), produced by the bacteria themselves^80,81,82–84^. The nanosilver-resistant strain (NAg^R^) upregulated the *pgaABCD* operon by ∼1.6-5.2-fold when exposed to the high NAg concentration (3 µg Ag/mL), relative to the cell-only samples (*p*-adj ≤ 0.01, **Figure 2A**). At the low nanoparticle exposures (0.5 x MIC), ∼2.2-3.4-fold upregulations were observed in NAg^R^, relative to WT. The operon encodes the membrane-bound protein complex PgaABCD, which synthesizes the exopolysaccharide poly-β-(1-6)-*N*-acetylglucosamine (PNAG), a major EPS constituent in *A. baumannii* biofilms^85^. The transcriptomic changes, however, were not seen in the WT strain upon silver exposure, relative to cell-only samples, hence defined as secondary defense mechanisms herein. Our phenotypic studies on biofilm growth supported these transcriptomic changes. As shown in **Figure 2C, D**, the resistant strain was associated with, not only a higher presence of surface-attached bacterial colonies, but also, a greater extent of EPS formation. Upon exposures to nanosilver (0.5 x MIC), a ∼2.4-2.5-fold more colonies (∼2130 µm^3^/µm^2^) and EPS formation (∼1545 µm^3^/µm^2^) were detected with the NAg^R^ biofilms, when compared to WT (∼850 and ∼654 µm^3^/µm^2^, respectively, *p* < 0.0001). The increased EPS formation, as reported in earlier studies, was likely to result from the *pgaABCD* upregulations seen with the resistant strain, relative to WT, conferring a greater antimicrobial defense^83,86^. The typically net negative-charged EPS has been indicated to adsorb and sequestrate heavy metals, including silver^87,88^. The presence of more colonies with the resistant strain biofilms could associate with the earlier described upregulations of the cell-to-surface adherence-associated outer membrane *ompA*, *ompW* and *carO*, as well as the membrane synthesis *lolA*, *blc* and capsule synthesis *otsA*, *otsB*, observed with the strain, relative to WT (at 0.5 x MIC nanoparticle exposure, **Figure 1A**), which is thought to enhance the bacterium colonization on surfaces. Further, the NAg^R^ strain was also found to upregulate *bap* that encodes one of the most conserved biofilm-associated proteins in *A. baumannii*, the Bap protein, which indeed, is also essential for cell-to-surface adherence of the bacterium, as well as for biofilm maturation^89^. A ∼6.1-fold increased *bap* expression was detected in NAg^R^, relative to WT, upon nanoparticle exposures (0.5 x MIC) (**Figure 2A**). The resistant strain also upregulated the conserved regulatory gene *bfmR*, part of the BfmR/S transcriptional stress response regulatory system, which, again, has been previously linked to biofilm formation in *A. baumannii*^90,91^. A ∼2.5-fold increased *bfmR* expression was observed in NAg^R^, relative to WT, upon nanoparticle exposures (0.5 x MIC). Studies have in fact reported BfmR/S transcriptional regulations of *bap*, *ompA*, as well as *otsA*, *otsB*^90,91^. The designation of the enhanced biofilm growth as secondary defense for nanosilver resistance, is in agreement with the non-observable increase, both in gene expressions (*pgaABCD*, *bap*, *bfmR*), and phenotypically, for the presence of surface-attached bacterial colonies and EPS formation with the WT strain, when exposed to the nanoparticle (0.5 x MIC), relative to the cell-only samples (**Figure 2A, C, D**).

**Figure 2.**
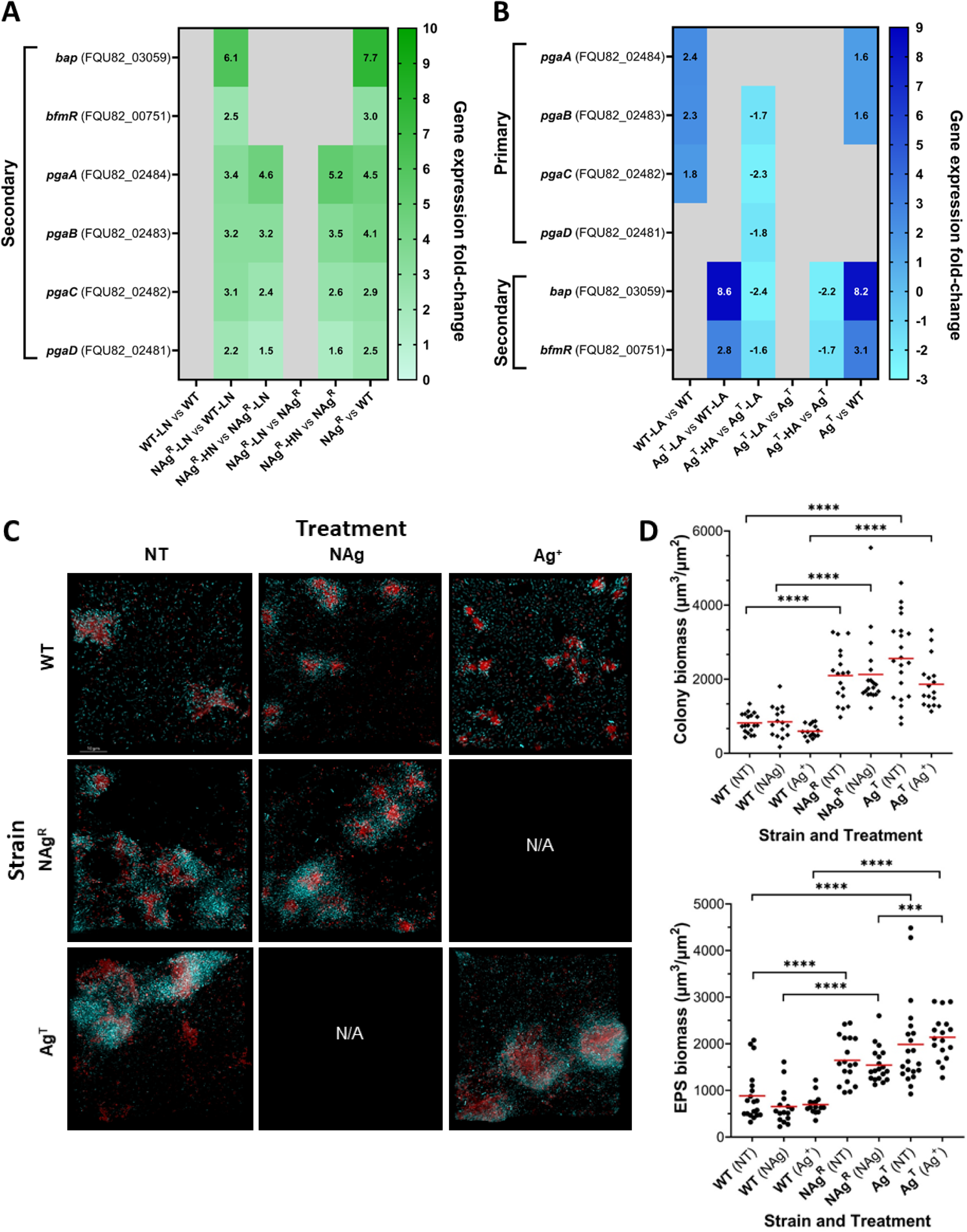
Changes in biofilm growth behavior in silver-adaptive *A. baumannii*. (**A, B**) Comparative analysis of chromosomal mRNA transcript levels of cell-to-surface adherence associated genes (*bap*, *bfmR*) and EPS synthesis genes (*pgaABCD*) in WT, NAg^R^ and Ag^T^ strains, upon exposures to the ‘low’ silver concentrations (0.5 µg Ag/mL for nanosilver (LN, 0.5 x MIC), 1 µg Ag/mL for ionic silver (LA, 0.5 x MIC)) and ‘high’ silver concentrations (3 µg Ag/mL for nanosilver (HN, 3 x MIC) and ionic silver (HA)). The differentially expressed genes (log_2_-fold change cut-off threshold ≥ 0.58 (≥ 1.5-fold change), *p*-adjusted ≤ 0.01, grey cells indicate statistically insignificant *p*-adj > 0.01 changes) were captured at 30 min of silver exposures in the exponential growth phase. Also shown are the physiological gene expression changes when in the absence of silver. (**C**) Fluorescent microscopy images of biofilms formed by WT and the silver-adaptive NAg^R^ and Ag^T^ strains when exposed to silver (0.5 x MIC nanosilver and/or ionic silver). Also shown is the respective cell-only no silver growth. Cyan color showed surface-attached bacterial colonies, while red color showed EPS mass, scale bar = 10 µm. (**D**) Quantitative analysis showed total colony (µm^3^/µm^2^) and EPS mass (µm^3^/µm^2^) of each strain under each respective silver exposures. Each data point represents calculated biomass surface coverages from individual biofilm images, with the horizontal bar showing the mean data from 15-20 images, per strain, per treatment. Statistical analysis (unpaired *t*-test with Welch’s correction) showed statistically significant differences in the bacterial colony and EPS abundance between strains and treatments (*** *p* < 0.001, **** *p* < 0.0001). The mRNA and phenotypic work were performed with a minimum of three biological replicates.

Further transcriptomic studies on the nanosilver-resistant strain showed upregulations of the EPS formation and the cell-to-surface adherence genes even when in the absence of silver. This indicates a stable biofilm-associated defense mechanism, which is also supported by phenotypic evidence. Upregulations of *pgaABCD* by ∼2.5-4.5-fold and *bap* by ∼7.7-fold were detected in NAg^R^, relative to WT (no silver, **Figure 2A**), and correspondingly, ∼1.9-fold higher extent of EPS formation (∼1645 µm^3^/µm^2^) and ∼2.6-fold more colony presence (∼2100 µm^3^/µm^2^) in NAg^R^, relative to WT (∼885 and ∼820 µm^3^/µm^2^, respectively, *p* < 0.0001, no silver, **Figure 2C, D**). Upregulation of the biofilm-associated stress response transcriptional regulator *bfmR* was also detected in the resistant strain, by ∼3.0-fold, relative to WT (no silver, **Figure 2A**). To recall, NAg^R^ also upregulated the cell-to-surface adherence-associated genes *ompA*, *ompW*, *carO*, *lolA*, *blc*, *otsA*, *otsB*, with no silver, relative to WT (**Figure 1A**).

These heightened biofilm growths were also seen in the case of ionic silver-tolerant strain (Ag^T^), again, both in the presence and absence of silver, the latter indicates a stable defense trait. Ag^T^ formed ∼3.2-fold more EPS (∼2205 µm^3^/µm^2^) with ∼2.9-fold more presence of colony biomass (∼1755 µm^3^/µm^2^) when exposed to ionic silver (0.5 x MIC), relative to WT (∼700 and ∼596 µm^3^/µm^2^, respectively, *p* < 0.0001, **Figure 2C, D**). Without silver, a ∼2.2-fold more EPS (∼1990 µm^3^/µm^2^) and ∼3.1-fold more colony (∼2560 µm^3^/µm^2^) were observed, relative to WT (∼885 and ∼820 µm^3^/µm^2^, respectively, *p* < 0.0001, **Figure 2C, D**). As with nanosilver, transcriptomic data also indicated a **secondary defense** for the biofilm growth behavior, with upregulations of *bap* by ∼8.6-fold and *bfmR* by ∼2.8-fold in Ag^T^, relative to WT, when exposed to ionic silver (0.5 x MIC) (**Figure 2B**). No statistically significant upregulations of these genes were detected in the WT strain upon ionic silver exposure, relative to the cell-only samples. Without silver, a ∼1.6-fold *pgaAB*, ∼8.2-fold *bap* and ∼3.1-fold *bfmR* upregulations were detected in Ag^T^, relative to WT (**Figure 2B**).

The observations highlighted changes in biofilm growth behavior in response to long-term silver exposures. While past inquiries have reported elevated biofilm growth in antibiotic-resistant bacteria, to the best of our knowledge, this is the first study to correlate stably heightened surface-attached colonization and EPS formation traits in a bacterium due to evolutionary silver adaptation^19,39^. With comparable transcriptomic and phenotypic trends, there were only slight differences in the biofilm growth characteristics between the silver-adaptive strains. A statistically significant (*p* < 0.001) higher extent of EPS formation was observed in the silver-exposed (0.5 x MIC) Ag^T^ biofilms, when compared to that of NAg^R^ (**Figure 2C, D**). Next, we described the evolved cellular defense responses to the well-established oxidative stress toxicity paradigm of silver.

### Oxidative stress defense and opportunistic silver efflux

As herein observed with the WT strain (0.5 x MIC, *p* < 0.0001, **Figure 3C, D**), bacterial exposures to nanosilver are known to generate cellular oxygen radicals, which, among others are the highly reactive one-electron reductant/oxidant superoxide and hydroxyl radicals that target biomolecules^24,29^. The nanosilver-resistant strain (NAg^R^) upregulated the superoxide dismutase *sodB* by ∼2.9-fold and catalases *katE*, *katG* by ∼14- and 1.8-fold, respectively, relative to WT when exposed to the low nanoparticle concentration (0.5 x MIC) (*p*-adj ≤ 0.01, **Figure 3A**). High nanoparticle concentration exposures (3 x MIC, 3 µg Ag/mL) saw further upregulation of *katG* by ∼2.3-fold in NAg^R^, relative to the low concentration exposures. The resistant strain also increased the expression of other catalase *katB*, by ∼3.4-fold, relative to the low concentration exposures. The enzyme superoxide dismutase SodB catalyses the dismutation of superoxide radical (O_2_^•-^) to molecular O_2_ and H_2_O_2._ The enzymes KatE (a hydroperoxidase, HPII), KatG (HPI), and KatB, then catalyze the reduction of H_2_O_2_ to O_2_ and H_2_O^92^. This reduction step would prevent cellular H_2_O_2_ conversion into other oxygen radicals, for instance, into the highly reactive hydroxyl radical (^•^OH), through the biologically common Fe(II)-mediated Fenton-type reaction of H_2_O_2_^24^. Studies have reported upregulations of *katE* and *katG* in *A. baumannii* under H_2_O_2_ oxidative stress, with the bacterium thought to express the latter catalase with heightened oxidative stress level^92–94^, as also seen herein with the high nanoparticle concentration exposures. It is also worth noting that *katE* was downregulated at the high concentration exposures, by ∼3.2-fold, relative to the low concentration exposures (**Figure 3A**). These upregulations of the ROS scavengers align with the less cellular ROS detected in NAg^R^, relative to WT, upon exposures to the nanoparticle (0.5 x MIC, *p* < 0.0001, **Figure 3C, E**). The resistant strain also increased the expression of *ohrB* by ∼1.9-fold, which encodes the organic hydroperoxide resistance protein OhrB which also helps control cellular H_2_O_2_ levels, but only at the low nanoparticle concentration exposures (0.5 x MIC), relative to WT (**Figure 3A**)^95,96^. Upregulations of these ROS scavengers were seen in NAg^R^, while absent in WT, hence referred to as secondary defense mechanisms. The WT only increased the expression of *acnA*, by ∼1.7-fold, with nanoparticle exposures (0.5 x MIC), relative to the cell-only samples (**Figure 3A**). The enzyme aconitate hydratase AcnA has been indicated to maintain the citric acid cycle during (or recovering from) cellular oxidative stress in *A. baumannii*^91,97,98,65^. The resistant strain also upregulated *acnA*, by ∼4.3-fold with the low nanoparticle concentration exposures (0.5 x MIC), relative to WT, and further, by ∼1.8-fold with the high nanoparticle concentration exposures (3 x MIC), relative to the low concentration exposures (**Figure 3**). Indicative of stable mechanisms, almost all ROS scavengers, *sodB*, *katE*, *katG*, *ohrB* and *acnA* (excluding *katB*), were innately upregulated in NAg^R^, without the presence of silver, by ∼1.7 – 30-fold, relative to the WT (**Figure 3A**).

**Figure 3.**
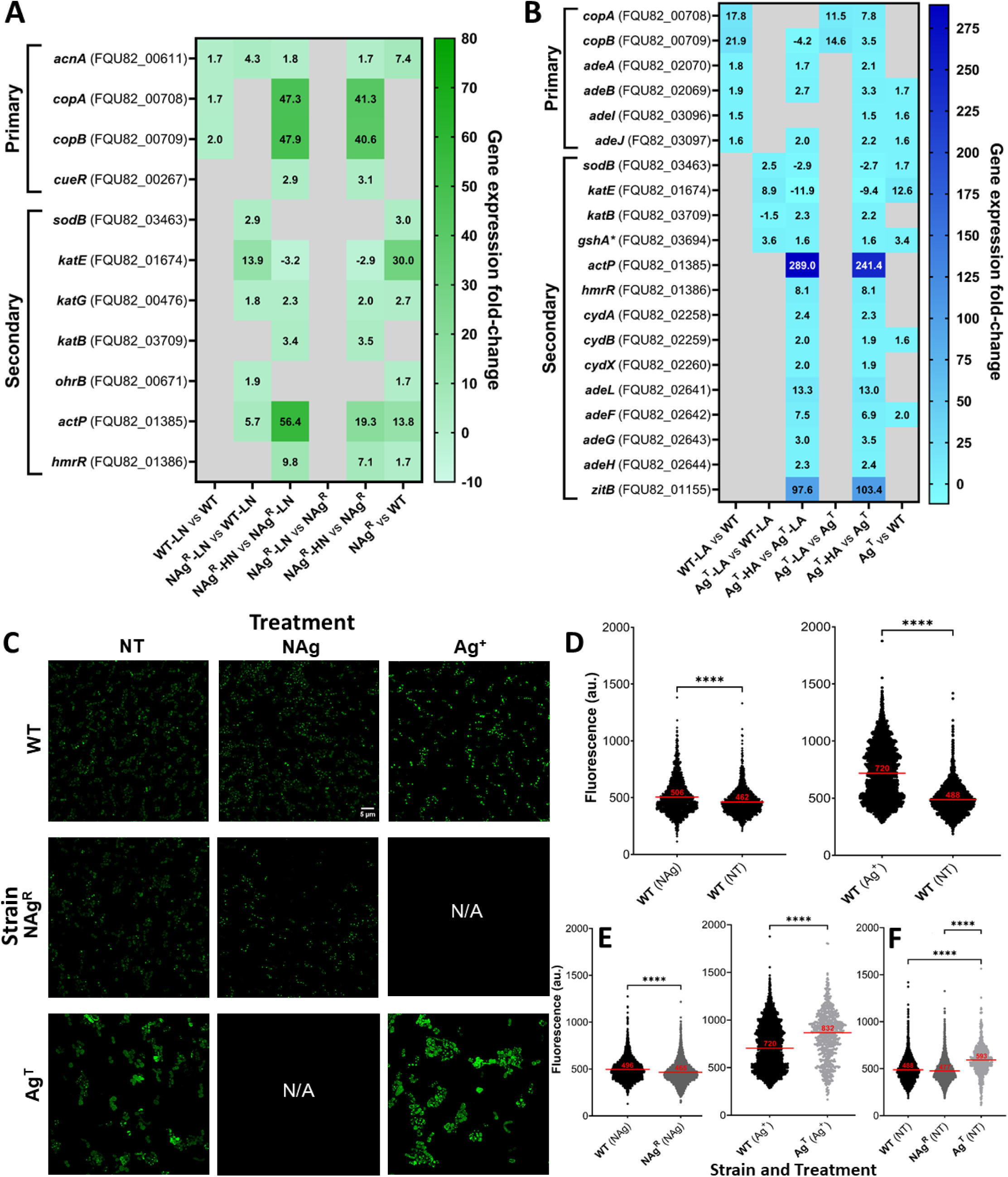
Oxidative stress management and efflux mechanisms in silver-adaptive *A. baumannii*. (**A**, **B**) Comparative analysis of chromosomal mRNA transcript levels of ROS scavenger and oxidative stress control (*sodB*, *katE*, *katG*, *katB*, *ohrR*, *acnA*) and metal efflux (*copA*, *copB*, *cueR*, *actP*, *hmrR*) genes in WT, NAg^R^ and Ag^T^ strains, upon exposures to the ‘low’ silver concentrations (0.5 µg Ag/mL for nanosilver (LN, 0.5x MIC), 1 µg Ag/mL for ionic silver (LA, 0.5 x MIC)) and ‘high’ silver concentrations (3 µg Ag/mL for nanosilver (HN, 3 x MIC) and ionic silver (HA)). Note the exclusive upregulations of the respiratory chain components (*cydA*, *cydB*, *cydX*) and efflux systems (*adeA*, *adeB*, *adeI*, *adeJ*, *adeL*, *adeF*, *adeG*, *adeH*) in Ag^T^, not seen in the case of nanosilver adaptation. The (*) denotes detection of single nucleotide mutation (in the oxidative stress control *gshA*). The differentially expressed genes (log_2_-fold change cut-off threshold ≥ 0.58 (≥ 1.5-fold change), *p*-adjusted ≤ 0.01, grey cells indicate statistically insignificant *p*-adj > 0.01 changes) were captured at 30 min of silver exposures in the exponential growth phase. Also shown are the physiological gene expression changes when in the absence of silver. (**C**) Fluorescence microscopy images of cellular ROS generation (green) in WT and the silver-adaptive NAg^R^ and Ag^T^ strains when exposed to silver (0.5 x MIC nanosilver and ionic silver, 30 min to 1 h). Also shown is the basal ROS detection in the respective cell-only no silver samples, scale bar = 5 µm. Quantitative analysis shows fluorescence intensity indicative of ROS levels in (**D**) silver-exposed WT to show silver-induced toxicity, and in (**E**) NAg^R^ and Ag^T^ to indicate ROS management in the adaptive strains. Also shown in (**F**) the physiological cellular ROS presence in WT, NAg^R^ and Ag^T^, without silver. Each data point represents single cell measurements, with the horizontal bar showing the mean data from 1000 – 4000 cells analyzed, per strain, per treatment. Statistical analysis (unpaired *t*-test with Welch’s correction) shows statistically significant differences in cellular ROS levels between strains and treatments (**** *p* < 0.0001). The mRNA and phenotypic work were performed with a minimum of three biological replicates.

Next, thought to serve as opportunistic silver defense mechanisms, the present work observed increased expressions of copper efflux genes, in WT, as well as in the nanosilver-resistant strain. A ∼1.7-2.0-fold upregulations of *copA* and *copB* were detected in WT, when exposed to nanosilver (0.5 x MIC), relative to the cell-only samples (**Figure 3A**), and by ∼47 – 48-fold in NAg^R^, the latter when exposed to the high nanoparticle concentration (3 x MIC), relative to those at low concentrations. CopA and CopB are inner membrane-bound P-type ATPase copper (Cu^+^) efflux pumps, which have been known to confer cross-resistance to silver ions in bacteria^99,100^. Earlier studies have reported upregulations of *copA* in response to silver exposures^76,77,101^. The upregulations, as also observed herein, have also been linked to less cellular ROS generation with metal exposures^100,102,76,77,101^. In *A. baumannii*, *copA* (and potentially *copB*) expressions are regulated by the HTH-type transcriptional regulator CueR^100,102,103^. Indeed, in conjunction with notable upregulations of *copA* (and *copB*) in the resistant strain, a ∼2.9-fold *cueR* upregulation was only seen in NAg^R^, with the high nanoparticle concentration exposures (3 x MIC), relative to those at lower concentrations (0.5 x MIC) (**Figure 3A**). The resistant strain also increased the expression of other copper (trans-membrane) P-type ATPase efflux pump *actP*^104,105^, by ∼56-fold, as well as, its transcriptional regulator *hmrR*^94^, by ∼9.8-fold, with the high nanoparticle concentration exposures (3 x MIC), relative to the low concentration (0.5 x MIC). No statistically significant changes in the expressions of *copA*, *copB* in Ag^T^, when in the absence of silver, relative to WT **(Figure 3A**), supporting the widely supported notion of heavy metal-induced upregulations of these efflux genes, in this case with silver^99,100,106^. As later described, physiological, without silver upregulations, were observed for efflux mechanisms, indeed, for non-heavy metal specific, multidrug pumps.

As in the case of nanosilver resistance, exposures to (ionic) silver also saw upregulations of the ROS scavenger superoxide dismutase and catalases, as well as the efflux pumps, in the ionic silver-tolerant strain (Ag^T^). Ag^T^ increased the expressions of *sodB* and *katE*, by ∼2.5-fold, ∼8.9-fold, respectively, relative to WT, when exposed to the low ionic silver concentration (0.5 x MIC, 1 µg Ag/mL) (**Figure 3B**). At the high silver concentration exposures (3 µg Ag/mL), Ag^T^ also upregulated *katB*, by ∼2.3-fold, relative to the low concentration exposures (0.5 x MIC). As with NAg^R^, Ag^T^ also downregulated *katE*, at the high concentration exposures, by ∼12-fold, relative to the low concentration exposures. The tolerant strain increased the expressions of *copA* and *copB*, with the low ionic silver concentration exposures (0.5 x MIC), by ∼12- and 15-fold, respectively, and by ∼7.8 and 3.5-fold, with the high silver concentration (3 µg Ag/mL), relative to the cell-only samples (**Figure 3B**). As also seen with NAg^R^, Ag^T^ also upregulated *actP* at substantial levels, by ∼290-fold, as well as *hmrR*, the transcriptional regulator, by ∼8.1-fold, with the high silver concentration exposures (3 µg Ag/mL), relative to the low concentration (0.5 x MIC). Notably, *copA* and *copB* were also upregulated in WT upon exposures to ionic silver (0.5 x MIC), by ∼18- and 22-fold, respectively, relative to the cell-only samples. The seemingly higher extent of *copA*, *copB* upregulations in WT than those in the ionic silver-adapted strain is in line with the non-observable upregulation of the transcriptional regulator *cueR* (in the latter strain). Interestingly, as later described, Ag^T^ upregulated various non-heavy metal specific, multidrug pump clusters, not seen with the nanosilver-resistant strain. In fact, despite the above-mentioned similarities in ROS scavenger and efflux expression changes, phenotypically, Ag^T^ was found to exhibit different cellular ROS generation profiles with (ionic) silver exposures, when compared to NAg^R^ with (nanoparticle) silver exposures. The tolerant strain, in fact, already displayed unusual cellular ROS profiles, inherently, in the absence of silver. As described in the next section, it is evident that the bacterium had also developed distinct defense mechanisms in response to ionic silver, particularly those involved in efflux mechanisms and oxidative stress control. This is thought to highlight, at least in part, the different adaptation characteristics of tolerance *versus* resistance, that evolved in response to the ionic *versus* nanoparticle silver prolonged exposures. A side note, when in the absence of silver, upregulations of *sodB*, *katE* were already detected in Ag^T^, by ∼1.7- and 13-fold, respectively, relative to WT, denoting a stable defense mechanism (Figure 3B). Again, as seen with NAg^R^, there were no statistically significant upregulations of *copA*, *copB* metal efflux genes in Ag^T^, in the absence of silver.

### Unique defense and physiological changes in ionic silver-tolerant strain

Recalling the earlier described overlapping (cell envelope and biofilm-associated) defense mechanisms to those of nanosilver adaptation, the present work found that the bacterium had also evolved distinct adaptation strategies in response to ionic silver prolonged exposures. The ionic silver-adapted strain has unique morphological and physiological traits. Firstly, the ionic silver-tolerant strain (Ag^T^) displayed irregular shapes with globules, in contrast to the well-defined rod shape of nanosilver-resistant strain (NAg^R^, as well as the WT) (**Figure 1C**). We hypothesize a correlation with the observed differences in membrane fluorescence intensity of the silver-adapted strains. A higher fluorescence (per µm of cell perimeter, *p* < 0.0001, **Figure 1C, D**) were detected in the (lipophilic) FM4-64-stained Ag^T^, relative to NAg^R^, which indicates a more compact membrane structure of the former. Earlier studies have reported morphological transformation in bacteria, including in *A. baumannii*, as a stress response to antibiotics, as well as silver, at sub-lethal levels^107–109^. The morphologically changed Ag^T^ indeed evolved from the WT following prolonged (ionic) silver exposure, initially at sub-lethal levels. One possible mechanism may involve OmpA. The outer membrane-embedded protein is anchored to periplasmic peptidoglycan, and its upregulation, as detected in Ag^T^, has been indicated to affect this outer membrane-peptidoglycan interaction, altering cell shape^110,111^. It remains unclear as to why we did not observe any morphological changes with NAg^R^, as OmpA upregulation was also seen in this train. Herein supported with more solid molecular evidence, physiologically, Ag^T^ was observed with higher cellular ROS presence, a reported characteristic of antimicrobial-tolerant strains^116,117^, relative to WT (*p* < 0.0001, as well as NAg^R^, *p* < 0.0001, **Figure 3C, F**), even when in the absence of silver. Tolerant bacterial populations have been indicated in many studies to enhance respiration, hence the elevated cellular ROS levels, as H_2_O_2_ is a natural by-product of respiration^116,117^. Note that this ROS build-up is independent of the silver-induced oxidative stress, also seen with exposures of WT strain to ionic silver (0.5 x MIC, 1 µg Ag/mL), relative to the cell-only samples (*p* < 0.0001, **Figure 3C, D**). In line with the elevated cellular ROS characteristics, the present work saw increased expressions of respiratory chain enzymes and efflux mechanisms, not seen in NAg^R^. A number of these distinct upregulations were also detected in Ag^T^ without silver, indicating they are stable mechanisms.

Ag^T^ upregulated *cydABX* by ∼2.0-2.4-fold, upon exposure to the high ionic silver concentration (3 µg Ag/mL), relative to the low concentration exposure (0.5 x MIC, 1 µg Ag/mL) (*p*-adj ≤ 0.01, **Figure 3B**). The gene cluster encodes the inner membrane-bound cytochrome *bd* oxidase complex CydABX, the terminal oxidase in the respiratory chain, which catalyses O_2_ reduction into water and drives the proton motive force for ATP generation^112^. Ionic silver (as well as ROS) has been indicated to target thiol groups in the membrane-bound respiratory enzymes, inhibiting their activities, in turn, disrupting the electron transport chain^24^. The *cydABX* upregulation is thought to compensate for the Ag^+^-targeting, hence maintaining the enhanced respiration characteristics in the tolerant strain. No statistically significant change in *cydABX* expression was observed in Ag^T^, relative to WT, with the low concentration exposures (0.5 x MIC), suggesting a higher-level ionic silver stress defense response. Further, the present work also detected upregulations of multidrug and heavy metal efflux systems in Ag^T^, not seen in the case of nanosilver, which could serve as additional mechanisms to protect from Ag^+^ targeting. The ionic silver-tolerant strain upregulated multidrug efflux pumps, which included the two known RND (resistance-nodulation-division) efflux gene clusters in *A. baumannii*, the *adeABC* and *adeIJK*. Ag^T^ increased the expressions of *adeA*, *adeB* by ∼1.7 and 2.7-fold, respectively, *adeJ* by ∼2.0-fold, upon exposures to the high ionic silver concentration (3 µg Ag/mL), relative to the low concentration exposures (0.5 x MIC, 1 µg Ag/mL) (**Figure 3B)**. In fact, upregulations of these genes were already observed in WT, at ∼1.5-1.9-fold for *adeA*, *adeB*, *adeI* and *adeJ*, in response to ionic silver exposures (0.5 x MIC), relative to the cell-only samples. Studies have indicated the roles of these efflux pumps in *A. baumannii* resistance to antibiotics, with emerging evidence on their roles in heavy metal resistance^113,114^. Also unique to ionic silver case, Ag^T^ upregulated the multidrug efflux *adeFGH* cluster^115^, by ∼2.3 – 7.5-fold, and indeed, as well as *adeL*, the transcriptional regulator, by ∼13-fold, with the high silver concentration (3 µg Ag/mL), relative to the low concentration exposures (0.5 x MIC). At more substantial level, Ag^T^ upregulated *zitB*, which encodes zinc(II) transporter ZitB, a member of the cation diffusion facilitator family, by ∼98-fold, with the high silver concentration (3 µg Ag/mL), relative to the low concentration exposures (0.5 x MIC). Upregulations of *zitB* have been observed in bacteria in response to toxic zinc exposures^116^. Interestingly, *zitB* over-expression, as herein observed, has been shown to be involved in defense responses to other heavy metals, such as iron and cobalt, suggesting that the protein may have non-specific functions, in this case, involving silver transport^117^. Our cellular ROS phenotypic observations with Ag^T^, when in the presence of silver, seem to be in line with the transcriptomic changes. Relative to WT, Ag^T^ was found with higher cellular ROS levels, when exposed to ionic silver (0.5 x MIC) (*p* < 0.0001, F**igure 3C****, E**). This contrasts with those earlier described with NAg^R^, whereby lower cellular ROS levels were detected in the nanosilver-resistant strain, relative to WT, with nanosilver exposures (0.5 x MIC). The phenotypic findings agree with the enhanced respiration characteristics hypothesized for the ionic silver-tolerant strain. A ∼1.6-2.0-fold stable upregulations of *cydB*, *adeB*, *adeI*, *adeJ* and *adeF* were detected in Ag^T^ (no silver, **Figure 3B**).

Thought to control cellular ROS levels, we found that Ag^T^ upregulated ROS scavengers, but, in general, involved less genes than those seen with NAg^R^. Like NAg^R^, the ionic silver-tolerant strain upregulated the superoxide dismutase *sodB* and catalases *katE*, *katB*, upon exposures to ionic silver (earlier described). The tolerant strain did not upregulate *katG* and *ohrB*, the latter also controls cellular H_2_O_2_ levels^95,96^, as well as *acnA*, for maintenance of citric acid cycle (during or post oxidative stress)^91,97,98,65^. An interesting single nucleotide mutation was observed in glutamate-cysteine ligase *gshA* in Ag^T^, which involves in the synthesis of the antioxidant glutathione (GSH)^118^. The substitutional (Gly39Arg) mutation was detected in 100% of the sequenced isolates^45^. Although still unclear at this stage, the mutation is thought to have a role in the observed upregulations of *gshA*, by ∼3.6-fold in Ag^T^, relative to WT, when exposed to the low ionic silver concentration (0.5 x MIC), and further, by ∼1.6-fold in Ag^T^, with the high silver concentration (3 µg Ag/mL), relative to the low concentration exposures (**Figure 3B**).

Herein, to summarize, we have established a working model for the adaptive defense mechanisms that evolved because of long-term nanosilver exposure (**Scheme 1**). The Gram-negative bacterium developed stable primary defenses, those of which were already exhibited in the WT, as well as secondary defenses, the latter only seen in the adapted strain. Cell surface defenses were activated with upregulations of outer membrane proteins as primary mechanisms, and membrane and capsule synthesis as secondary mechanisms. The phenotypically indicated increase in membrane integrity of the nanosilver-resistant strain (NAg^R^) is thought to also enhance the cell-to-surface adherence behavior of the bacterium, which could be associated with the observed increase in the extent of protective biofilm growth. The work saw more surface-attached growth of bacterial colonies, as well as higher abundance of the EPS matrix shields, with the resistant bacterium also upregulating more cell-to-surface adherence genes, as well as EPS synthesis genes, as secondary mechanisms. Finally, as anticipated, the bacterium had also evolved defenses to adapt to the known oxidative stress-mediated toxicity paradigm of nanosilver. ROS scavengers were upregulated as secondary mechanisms in the resistant bacterium, along with the supposedly opportunistic silver efflux mechanisms, the latter already seen in WT strain. The bacterium evolved different adaptation characteristics in response to long-term ionic silver exposures. The slower-to-kill ionic silver-tolerant strain (Ag^T^) had developed a generally comparable primary and secondary cell surface defenses, with similar upregulation patterns for outer membrane proteins and capsule synthesis genes, respectively, as with NAg^R^. The tolerant strain also heightened its biofilm growth behavior, with more surface-attached colony growth, which, as with NAg^R^, is thought to associate with the phenotypically indicated increase in membrane integrity. The tolerant strain, however, was found to form more EPS when compared to NAg^R^, although the reasons for this remain unclear. Indeed, Ag^T^ is physiologically different to NAg^R^. The tolerant strain was associated with higher cellular ROS profiles, a known antimicrobial-tolerant trait, being linked to enhanced respiration activities. Not seen in NAg^R^, the tolerant strain upregulated the respiratory chain terminal oxidase cytochrome, as well as several multidrug and heavy metal efflux systems, thought to serve as defense mechanisms. More specifically, to replenish the respiratory chain component from the known Ag^+^ targeting, and to fend off Ag^+^ intake, respectively. Ag^T^ also exhibited different oxidative stress control mechanisms, when compared to NAg^R^, overall upregulating less ROS scavengers or ROS control-associated genes. At this stage, the exact reasons for the differing adaptation characteristics that evolved from long-term exposure to the nanoparticle, as opposed to the ionic form of silver, remain unclear. Apart from their unique silver antimicrobial characteristics, the stable morphological transformation seen with Ag^T^ could perhaps give us further clues for the different adaptation responses. Studies have suggested that bacteria change shape to either increase their surface-to-volume ratio, hence enhancing the rate of nutrient uptake/antimicrobial efflux, or to decrease the ratio, therefore limiting antimicrobial influx^119,120^. As shown in **Figure 1C**, the latter seems to apply for the ionic silver-tolerant strain, with larger cell size (relative to WT, and NAg^R^), possibly to reduce intake of silver ions.

**Scheme 1.**
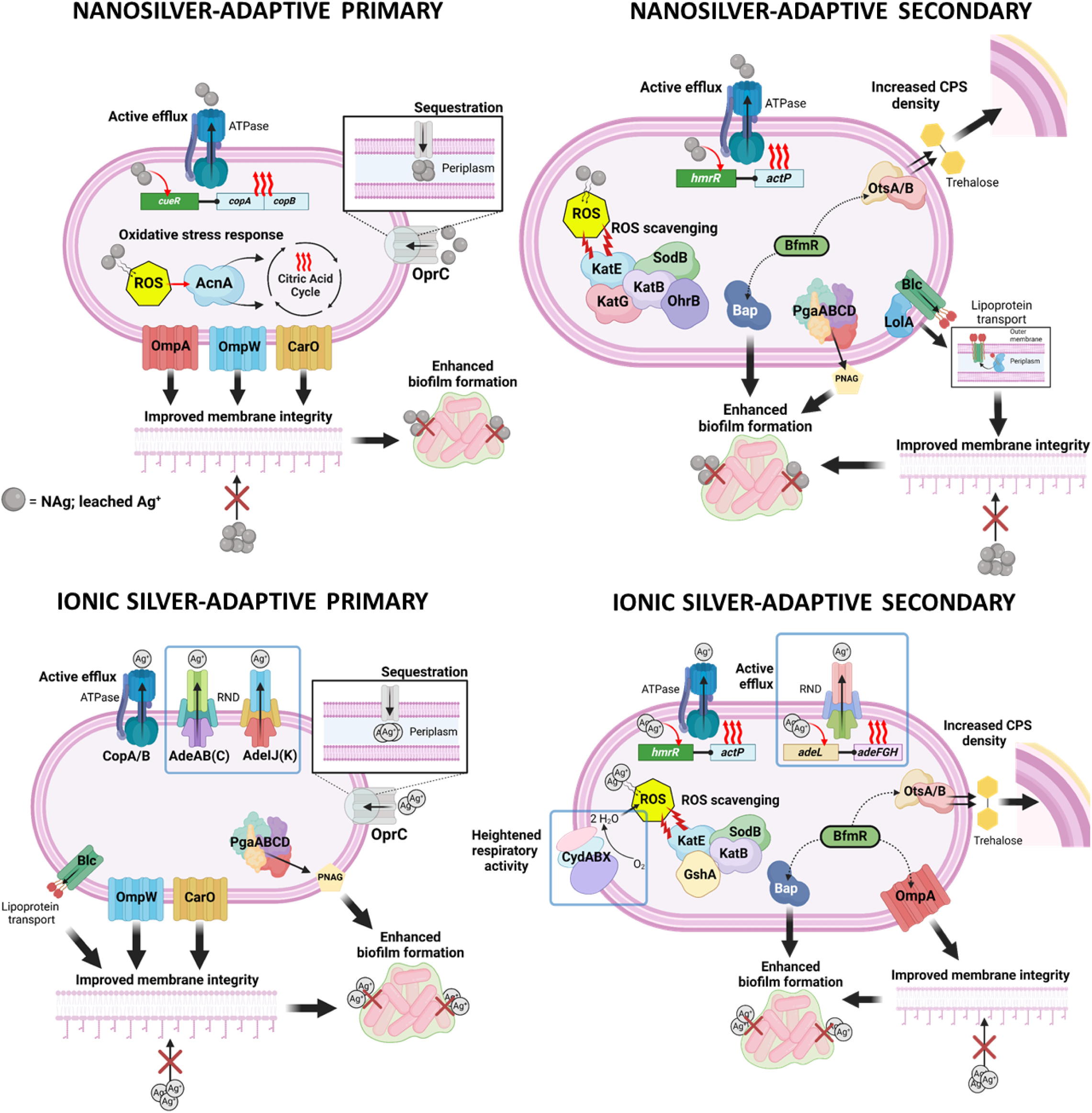
Working models of silver-adpated defense mechanisms in the Gram-negative bacterium *A. baumannii*. The silver-adaptive strains exhibited both primary and secondary mechanisms, with the wild-type only displaying primary mechanisms. Also shown (in blue frames) are the unique mechanisms that evolved in the case of ionic silver adaptation. Created in [BioRender](https://www.biorender.com/).

Finally, the study also revealed a plasmid-oriented silver stress response mechanism. A transcriptomic analysis of native plasmids in the bacterium (p1 of 7655 bp and p2 of 9540 bp)^49^ showed upregulations of type II toxin/antitoxin (TA) *higBA* (present in both p1 and p2), by ∼2.8-4.5-fold and ∼1.7-3.5-fold in NAg^R^ and Ag^T^ strains, respectively, when exposed to the high concentration of their respective silver agents (3 µg Ag/mL), relative to the low silver concentration (0.5 x MIC) exposures, and the cell-only samples (see **Figures S9** and **S10**). With respect to silver toxicity, studies have indicated the roles of the type II TA in modulating stress responses in bacteria, including in response to oxygen radical-targeting^121,122^. While yet to be shown in *A. baumannii*, HigBA has been suggested to play a role in DNA repair. Studies with *Enterobacteriaceae* plasmids have highlighted the presence of LexA binding sites in the *higBA* promoter regions. LexA works together with RecA as an SOS response system, whereby binding of RecA to single strand (damaged) DNA will induce the self-release of LexA, which in this case, would lead to upregulation of *higBA*^123^. Both NAg^R^ and Ag^T^ upregulated *recA* by ∼1.9- and 2.3-fold, respectively, at the high silver concentration, relative to the low concentration (0.5 x MIC) exposures (**Table S4**). HigBA has also been associated with enhanced biofilm growth in response to stressors^121,123^, the latter also manifested with silver-adapted defense.

## Conclusion

The molecular work described the adaptive defense mechanisms of a Gram-negative pathogen, which evolved in response to long-term exposures to nanosilver, and how these are different to those developed in response to ionic silver. The evolutionary adaptation saw the resultant ‘harder-to-kill’ nanosilver-resistant bacterium upregulating, not only its ROS scavenger and heavy metal efflux mechanisms, which was anticipated, but also, its cell surface defense systems. More specifically, these included various outer membrane proteins, along with membrane and capsule synthesis genes, for the latter. Interestingly, the bacterium also produced more biofilms, exhibiting a higher presence of surface-attached colonies, which was most likely associated with the indicated increase in membrane density. The bacterium also expressed a greater abundance of the protective EPS biofilm matrix. These stable defense traits manifested as primary and secondary mechanisms, the latter were observed only in the silver-adaptive strains, while absent in the wild-type. Although sharing similarities in cell surface and biofilm defense traits as with the nanosilver adaptation case, the ‘slower-to-kill’ ionic silver-tolerant strain further upregulated its respiratory chain enzymes and multidrug efflux systems. These exclusive mechanisms align with emerging scholarly evidence of enhanced respiration characteristics in antimicrobial-tolerant bacteria, in this case, to replenish the known cellular targets of Ag^+^, and to minimize cellular intake of the ions, respectively. The findings highlight these unique antimicrobial characteristics, and therefore potentially distinct bacterial adaptation responses to the nanoparticle *versus* ionic forms of silver. The elucidation of the adaptation phenomena can inform strategies to control the risks of long-term silver exposures. The identified tiered defense mechanisms at mRNA levels can serve as molecular targets to overcome the resistance phenomena, to ensure successful long-term use of these important nano-antimicrobials.

## Acknowledgements

We would like to acknowledge the Australian Research Council for the research funding (ARC Discovery Projects DP180100474, DP220101819, and DP220101143) and the UTS Science PhD Scholarship for O.M; B.S. is supported by the ARC Future Fellowship (FT230100062). We would also like to acknowledge the technical assistance from Dr Igor Makunin (Queensland Cyber Infrastructure Foundation, QCIF, Australia) for the RNAseq analysis, Associate Professor Louise Cole and Dr Amy Bottomley (Microbial Imaging Facility, MIF, University of Technology Sydney, Australia) for their support in the phenotypic imaging studies, as well as Scientia Professor Rose Amal and Dr Emma Lovell (the University of New South Wales, Australia) for the silver nanoparticles.

## Supporting Information

**Table S1.** Working concentrations of NAg and Ag^+^ for WT, NAg^R^, and Ag^T^ *A. baumannii* ATCC 19606 RNA and phenotypic studies.

**DOI: 10.6084/m9.figshare.25955890**

**Table S2.** Gene ontology enrichment analysis of statistically significant differentially expressed genes (DEGs) mapped to functional biological pathways identified in the RNA-seq comparisons of WT and NAg^R^.

**DOI: 10.6084/m9.figshare.25956058**

**Table S3.** Gene ontology enrichment analysis of statistically significant differentially expressed genes (DEGs) mapped to functional biological pathways identified in the RNA-seq comparisons of WT and Ag^T^.

**DOI: 10.6084/m9.figshare.26010916**

**Table S4.** DEGs (upregulated and downregulated genes) identified in WT and NAg^R^ exposed to low (0.5 x MIC, 0.5 µg Ag/mL) and/or high concentrations (3 µg Ag /mL) of NAg.

**DOI: 10.6084/m9.figshare.26011252**

**Table S5.** DEGs (upregulated and downregulated genes) identified in WT and Ag^T^ exposed to low (0.5 x MIC, 1 µg/mL) and/or high concentrations (3 µg Ag/mL) of Ag^+^.

**DOI: 10.6084/m9.figshare.26011270**

**Figure S1.** Schematic workflow summary of RNA-seq analysis.

**DOI: 10.6084/m9.figshare.25955929**

**Figure S2.** Principal component analysis (PCA) of RNA studies.

**DOI: 10.6084/m9.figshare.26011012**

**Figure S3.** Comparative growth rates of WT when exposed to low NAg concentration (0.5 x MIC, 0.5 µg Ag/mL) and the cell-only growth (no silver).

**DOI: 10.6084/m9.figshare.26011282**

**Figure S4.** Comparative growth rates of WT and NAg^R^ when exposed to low (0.5 µg Ag/mL) and high (3 µg Ag/mL) NAg concentrations.

**DOI: 10.6084/m9.figshare.26011540**

**Figure S5.** Comparative growth rates of cell-only (no silver) WT and NAg^R^.

**DOI: 10.6084/m9.figshare.26011552**

**Figure S6.** Comparative growth rates of WT when exposed to low Ag^+^ concentration (0.5 x MIC, 1 Ag µg/mL) and the cell-only growth (no silver).

**DOI: 10.6084/m9.figshare.26011558**

**Figure S7.** Comparative growth rates of WT and Ag^T^ when exposed to low (1 µg Ag/mL) and high (3 µg Ag/mL) Ag^+^ concentrations.

**DOI: 10.6084/m9.figshare.26011567**

**Figure S8.** Comparative growth rates of cell-only (no silver) WT and Ag^T^.

**DOI: 10.6084/m9.figshare.26011570**

**Figure S9.** Differentially expressed genes (DEGs) identified in plasmid p1 of WT, NAg^R^, and Ag^T^.

**DOI: 10.6084/m9.figshare.26011591**

**Figure S10.** Differentially expressed genes (DEGs) identified in plasmid p2 of WT, NAg^R^, and Ag^T^.

**DOI: 10.6084/m9.figshare.26011609**

